# Biofilm formation and toxin production provide a fitness advantage in mixed colonies of natural yeast isolates

**DOI:** 10.1101/178244

**Authors:** Bernadette M. Deschaine, Angela R. Heysel, Adam Lenhart, Helen A. Murphy

**Affiliations:** Department of Biology, The College of William and Mary, Williamsburg, Virginia

**Keywords:** colony morphology, biofilm, sociomicrobiology, Saccharomyces cerevisiae, K2 toxin, K28 toxin

## Abstract

Microbes exist in complex communities and can engage in social interactions ranging from cooperation to warfare. Biofilms are structured, cooperative microbial communities; they are pervasive and ancient, representing the first fossilized life. Like all cooperative communities, biofilms are susceptible to invasion by selfish individuals who benefit from cooperation, but do not contribute. The ubiquity of biofilms therefore poses a challenge to evolutionary theory. One hypothesis for biofilm stability is spatial structure: patches of related cooperative cells are able to outcompete unrelated cells. These dynamics have been explored computationally and in bacterial systems; however, their relevance to eukaryotic microbes remains an open question. Here, we investigate the interactions of environmental yeast isolates with different social phenotypes. Our results show that biofilm strains spatially exclude non-biofilm strains, and that biofilm spatial structure confers a consistent and robust fitness advantage in direct competition. We also find that biofilms protect against killer toxin, a warfare phenotype. During biofilm formation, cells are susceptible to toxin from nearby competitors; however, increased spatial use by biofilms provides an escape from toxin-producers. Our results suggest that yeast biofilms represent a competitive strategy, and that principles elucidated for the evolution and stability of bacterial biofilms may apply to eukaryotes.

## Background

For most of the time since the discovery of microorganisms, microbes were thought to live a relatively solitary existence, interacting mostly with their environment. It has now become clear that social interactions between microbes, both within and between species, are abundant and extremely important. Such interactions can include cooperation, competition, synchronization, and even chemical warfare [1]. Biofilms are cooperative microbial communities composed of one or multiple species, anchored to a surface, and protected from environmental hazards by a secreted extracellular matrix [2, 3]. They are found throughout the natural and manmade environment, wherever microbes are found. The oldest fossils on earth are microbial mats; thus, it appears that there have been biofilms since microbes first evolved [4].

Biofilms require individuals to produce goods, such as components of the extracellular matrix, that can be used by all members. Like all cooperative communities, they are susceptible to “ cheaters” who do not produce the public goods, yet benefit from them [5-8]. Despite their vulnerability to individual cheaters, biofilms are ubiquitous. The leading hypothesis for the stability of biofilm communities is the spatial structure: competition, cooperation, and passive processes like clonal growth can generate patches of related cooperative cells able to outcompete unrelated cells (e.g., [9-18], recently reviewed in detail in ref. [19]). Aside from acting as a public good, the production of substances that facilitate cell-to-cell and cell-to-surface adherence can be a competitive cooperative strategy that allows lineages increased access to space and nutrients [10, 20-22], and can even work to exclude non-producers from the community [23]. Recent work in the bacterium *Pseudomonas aeruginosa* demonstrated that when multiple strains were grown together, biofilm formation increased, and single strains often dominated the competition [24].

Another type of competitive strategy in microbial communities is warfare, which takes the form of microbial toxins and antibiotics [25, 26]. Under certain conditions, warfare-producing and sensitive lineages can coexist within an expanding spatially structured community [27-31]. It has also recently been demonstrated that in a dense, well-mixed community, a warfare phenotype can generate spatial segregation of producing and sensitive lineages[32]. The interaction between microbes producing warfare phenotypes and microbes producing biofilms is not yet entirely clear. The same study that investigated multi-strain *P. aeruginosa* communities also found that the production of antibiotics by competitors increased biofilm formation [24]. This suggests that biofilms may serve to protect from warfare phenotypes.

Most research on microbial social evolution has been conducted in bacterial systems[1, 5, 19, 33]. However, the complexity of eukaryotic cells and the potential differences between bacterial and fungal biofilms [34] leave open the possibility that the social dynamics may be quite different in eukaryotic microbes. Furthermore, the relevance of fungal biofilms to public health [35] suggests that understanding the social and evolutionary dynamics within a fungal model is of increasing importance.

### Saccharomyces cerevisiae Social Phenotypes

Cells of the model yeast, *Saccharomyces cerevisiae*, can adhere to each other and various surfaces, forming mats, floccs, and complex colonies [36-40], and can also engage in warfare through toxins[26]. As researchers have amassed a global collection of isolates [41-45], it has become clear that these social phenotypes are common [46-48], making *S. cerevisiae* an ideal model to study fungal biofilms[49, 50] and investigate questions related to eukaryotic sociomicrobiology.

Yeast colonies with complex colony morphology can form on solid agar surfaces [48]. These “ fluffy” colonies [36] resemble the wrinkly colonies of the bacterial biofilm models *P. aeruginosa* and *Bacillus subtilis* and have all the hallmarks of fungal biofilms[34]: an extracellular matrix facilitating nutrient flow and water retention[36, 51]; expression of drug efflux pumps; and velcro-like structures attaching cells to one another [52] encoded by an adhesin gene, *FLO11*[53]. When grown as single-strain colonies[54] or mats[55], strains forming biofilms have been shown to spread and occupy space more quickly than non-biofilm-forming (smooth) strains; however, smooth colonies have a greater cell density[51]. Thus, cell counts, rather than colony size, should be used to test the fitness effects of biofilm formation. While simple smooth *S. cerevisae* colonies have been used to explore spatially expanding mixed populations (e.g., [13, 16, 18, 56]), and one study has generated mixed *FLO11* and *flo11* colonies from a single lab background [57], to our knowledge, the evolutionary dynamics of multi-strain biofilm communities have not been explored.

Yeast killer toxins are secreted proteins that function in inter-strain competition: secreting cells are protected, while nearby sensitive cells are killed [26]. Killer toxins are encoded by cytoplasmically inherited double-stranded RNA (dsRNA) viruses; they replicate with the aid of dsRNA helper viruses, which encode the viral capsid protein and RNA-dependent RNA polymerase[26]. Several different yeast killer toxins with different modes of action have been identified [58-60]. Toxins occur widely in natural populations of *Saccharomyces* yeasts, with toxin production detected in ∼10% of strains surveyed from publicly available *S. cerevisiae* and *S. paradoxus* collections[47].

The research presented here focuses on two killer toxins, K2 and K28, and their effects on biofilm-producing strains. K2 [61] is the killer toxin most commonly found in vineyard ecosystems[47]. It acts quickly to induce membrane permeability and reduce intracellular ATP levels in sensitive cells, but the details of its mode of action remain unknown[58]. K28 was first identified in an *S. cerevisiae* wine strain [62]. It is taken up by sensitive cells and interferes with proteins essential for cell-cycle control, fatally blocking DNA synthesis [63].

### This Study

We sought to test the generality of the predictions of microbial social evolution theory by examining two social phenotypes in a eukaryotic system. We first explored whether clonal growth and spatial structure provide fitness benefits in yeast. Based on computational and experimental work with bacterial species (e.g., [10, 20, 23, 64]), we hypothesized that biofilm-formers would outcompete non-biofilm-formers for both space and resources. Next, we tested whether biofilm production protected cells from killer toxin, an area of microbial social evolution with far less background theoretical and experimental research.

Our experiments used environmental isolates in order to understand how natural social phenotypes may interact. *S. cerevisiae* is found in a variety of ecological niches (woodlands, vineyards, industrial, agricultural, and clinical settings [41-45]), and insects have been shown to transport the yeast and to increase outcrossing rates[65-68]. This suggests that different genetic backgrounds likely interact in nature and may directly compete with one another. We therefore directly and indirectly competed isolates of *S. cerevisiae* in spatially structured communities, using cell counts to determine the fitness effects of biofilm formation and toxin production. Our experimental results demonstrated a consistent and robust fitness benefit to biofilm formation in direct competition, and suggest that spatial use may provide a way to escape from toxin-producing competitors.

## Methods

### Strains

*S. cerevisiae* from publically available [41, 42] and personal collections were screened to identify strains of interest [48] (table S1). Two biofilm strains-YJM311 (clinical) [69], YJM224 (distillery yeast), and three smooth strains-YJM981 (clinical) [70], SK1 (lab/soil) [41], YPS681 (woodland) [71], were selected for fitness assays. The diploid isolates were transformed [72] with a cassette that targeted the terminal region of the highly expressed *PGK1* gene (table S2) and contained: (1) either mCherry or GFP, and (2) antibiotic resistance through KanMX [73], NatMX or HghMX [74]. Plasmids pFA6a-GFP-KanMX6 [75] and pBS34-mCherry-KanMX6 [76] were used as template for PCR (Yeast Resource Center, University of Washington); the antibiotic resistance was subsequently switched via transformation for some strains. Strains with killer toxin virus, K28 (MS300b) [77] and K2 (29-06) [47] (generously provided by D. Wloch-Salamon) were used for assays with toxin activity [47]. K28 is active in acidic conditions (∼3.0-5.0 pH), with optimum activity at a pH of ∼4.7 at temperatures between 20-30°C[78, 79], while K2 is active in the acidic pH range of 2.5-5.0 at temperatures between 20-25°C[78].

### Media

Strains were grown in YPD (1% yeast extract, 2% peptone, 2% dextrose) or low dextrose (LD) YPD (0.1% dextrose); solid media contained 2% agar. When appropriate, media was supplemented with 150 µg/ml G418, 75 µg/ml CloNat, or 300 µg/ml hygromycin B. Toxin assays were performed on 1.5% agar YPD and LD-YPD plates supplemented with citric acid to adjust the pH to 4.5 (∼3 mg)[47, 78], and with methylene blue, which stains dead yeast cells allowing visualization of the toxin activity[80].

### Fitness Assays

*Start*: 2 µL of overnight culture was added to 198 µL of water in wells of a non-treated 96-well plate. For mixed colonies, for a 1:1 ratio, 1µL of each strain was added; for a 1:9 ratio, 2 µL of an appropriately mixed culture was added. Three replicates were made for each strain and mix of strains; in a given 96-well plate, only 15 wells were used, such that each experimental well was surrounded by empty wells. Cultures were then pinned onto YPD and LD-YPD OmniTrays (Nunc 264728) using a 96-pin multi-blot replicator (V&P Scientific no. VP408FP6). For assays with toxin-producing strains, cultures were also pinned onto low-pH YPD, and low-pH LD-YPD. Initial cell counts were made in one of two ways: (1) plating culture from the wells, followed by replica-plating to appropriate antibiotic plates or viewing colonies under a fluorescence stereoscope for mixed colonies, or (2) imaging a sample of the culture with a hemacytometer and differentiating strains from mixed cultures with fluorescence markers. Colonies were started with ∼500-1000 cells, as previous work has shown that starting with a low density can itself generate spatial segregation[17].

*Growth:* For assays with non-toxin strains, YPD plates were incubated for 3 days and LD-YPD for 5 days, both at 30°C. With toxin-producing strains, all plates were incubated at room temperature for approximately 6-7 days [78]. For mechanical disruption, when colony growth was evident, a sterile pin was used to swirl colonies once per day. Fluorescent and/or light images were taken of each colony (Zeiss SteREO Discovery.V12 and Nikon D3200 camera) and processed in Fiji [81].

*End:* To maximize recovery of cells for sampling, entire colonies were removed from the plates using a metal cylinder with attached rubber bulb (figure S1). The agar plugs were suspended in either 2.5 mL water or 15% glycerol (when stored for later processing). The cylinder was sterilized via ethanol and flaming between plugs. In order to separate cells adhering to agar and/or other cells, several sterile 3.5 mm glass beads were added to each tube and gently sonicated (UP200St sonicator with VialTweeter Sonotrode). Final cell counts were made from the processed colonies via plating or imaging with a hemacytometer, as described above.

Since there are multiple sources of experimental variation in this assay, all natural strain combinations were assayed independently by two different researchers (A.H. and B.D.) using slide counts. All natural strain combinations were also assayed by plate counts to verify that cells survived colony processing. Toxin assays were performed using plate counts.

### Competitions in Liquid Assays

Competitions were initiated with overnight cultures grown in the medium in which the competition would occur; 10ul of a single strain, or 10ul of a 1:1 (by volume) mix of two strains were inoculated into 10mL of YPD or LD. Initial counts of each culture and master mix were made using a hemacytometer. Cultures were grown at 30°C with shaking; 10ul was serially transferred every 24 hours for either 2 or 3 cycles. After 48-72h from the start, final cell counts were made with a hemacytometer.

### Statistical Analysis

The data were analyzed in JMP v11.2.0 using a least-squares regression model with biofilm strain and treatment (medium + single vs. mixed community) classified as fixed effects, and assay type (plate counts vs. slide counts) and researcher (A.H. and B.D.) as random effects. The data were analyzed two ways, with the response variable as either the relative change of the biofilm strain or the change in the frequency (slope) of the biofilm strain.

## Results

To determine the fitness effect of biofilm formation during competition between unrelated genetic backgrounds, yeast strains were grown in spatially structured communities on agar plates. The experiments focused on two biofilm strains with distinct colony morphologies, which were competed against three non-biofilm strains, and then against two toxin-producing strains. Pairs of biofilm and non-biofilm strains were competed against one another in both homogenous and mixed communities. Biofilm formation is induced in carbon-limited conditions[48]; strain pairs were assayed with and without biofilm induction by adjusting the amount of glucose in the medium. Both K2 and K28 toxins are active only in acidic conditions; strain pairs were assayed with and without toxin activity by adjusting pH.

In contrast to microbial competitions performed in liquid, spatially structured colonies do not meet the assumptions of traditional fitness calculations, specifically the requirement of a well-mixed population[82]. Instead, growth is mostly limited to the front at the leading edge of the colony[9]. Therefore, the change in the proportion of the biofilm strain was used as a proxy for fitness. Simply by chance, a strain in a mixed colony could "win" a competition by reaching and monopolizing the edge of the colony, and thus greatly increase its proportion of the population. However, by averaging over replicate colonies, the effect of chance is minimized and the competitive ability of a given strain should become clear. Each of our assays included three replicates of each strain or mixed pair, and assays were performed multiple times.

### Competitions between biofilm and non-biofilm strains in spatially structured communities

Our results demonstrate a consistent and robust fitness advantage to biofilm formation in direct competition (figure 1). With relative change in the frequency of the biofilm strain as the response variable, treatment and biofilm strain were significant (p<0.0001, p=0.0232, respectively). The LD-mixed community treatment was significantly greater than the other three treatments (Tukey's HSD, ɑ=0.05), and overall, YJM311 was significantly more fit than YJM224 in competitions with other natural strains. Assay type (slide counts vs. plate counts) and researcher performing the experiment contributed 1% and 12% of the variation, respectively. This analysis was repeated using the difference in proportion of the biofilm strain as a response variable and produced qualitatively similar results, although the identity of the biofilm strain was not significant (figure S3; table S3).

**Figure 1:**
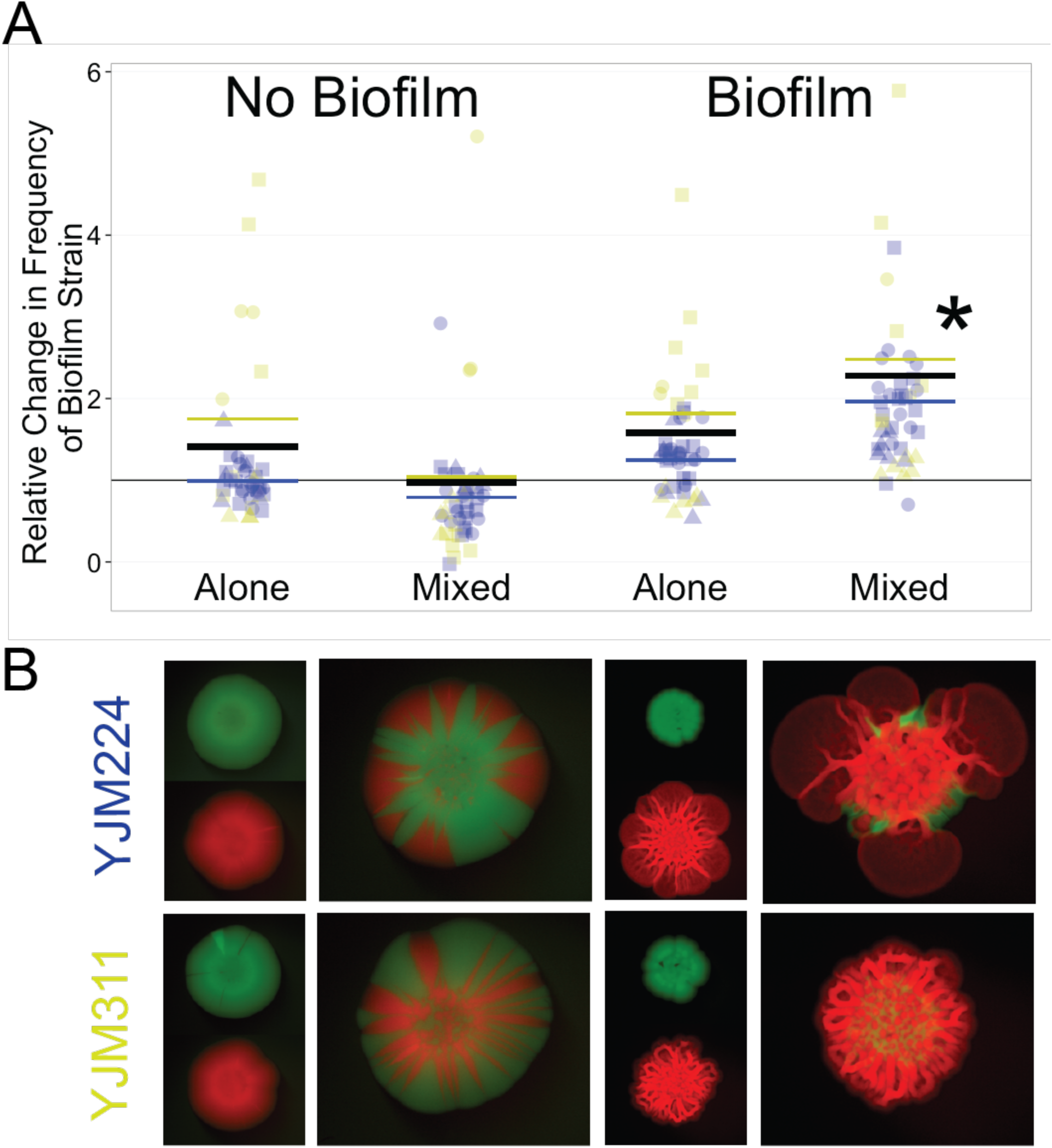
Fitness Effects of Biofilm Formation. Biofilm-forming strains were competed against non-biofilm-forming strains in pure and mixed colonies, with and without inducing biofilm formation. For the pure colony treatment, colonies were paired at random and the frequency of each strain was estimated. A total of 240 colonies were assayed. **(A)** Colors correspond to the biofilm-forming strains listed in (B), YJM224 and YJM311; shapes correspond to identity of non-biofilm-forming strains: circle-SK1, triangle-YJM981, square-YPS681; * indicates significance at p< 0.0001. Black lines represent overall mean for a treatment; colored lines represent biofilm-strain mean. **(B)** Representative images of the experimental treatments, as labeled in (A). Mixed colonies are to scale relative to one another; pure colonies are to scale relative to one another, but are scaled to half the size of the mixed colonies. Each row represents a single strain combination.

Inspection of figure 1*a* shows that with the exception of a single colony, the biofilm strain increased in frequency relative to the non-biofilm strain in every mixed colony grown on LD. In contrast, in mixed communities grown on YPD, the biofilm strain had similar or decreased fitness. Growth was also compared between homogenous, single-strain colonies of biofilm and non-biofilm strains. The biofilm strains had similar fitness to non-biofilm strains in both LD and YPD conditions. This suggests that the relative increase of the biofilm strains in mixed colonies is not simply due to faster growth in low dextrose conditions, but rather a competitive ability provided by biofilm formation. Based on the gross morphology of the mixed colonies (figure 1*b* and S2), we hypothesized that the advantage was due to the spatial structure of the community, specifically the biofilm strain's ability to monopolize the leading edge of the colony. Importantly, biofilm strains appear to be able to spatially exclude non-biofilm strains.

To test this hypothesis, two further experiments were performed based on the following logic: If the ability to increase in frequency were due to reaching and monopolizing the edge of the colony quickly, the biofilm strain's competitiveness could be hampered by either: (1) starting with a low frequency of the biofilm strain, thus lowering the probability of reaching the edge first, or (2) mechanically disrupting the spatial structure of the community during growth. First, the assay was repeated with the mixed colony inoculum prepared at a 1:9 biofilm to non-biofilm strain ratio, rather than 1:1. In the original assay, due to the inherent variation of the procedure, differing growth rates, and experimental error, the starting ratio of the colonies ranged from 0.07 to 0.9 (figure S3). The 1:9 assay was intended to more systematically test competitive ability in low-frequency conditions, and used a smaller subset of strain combinations. The results showed an even more dramatic competitive advantage to forming a biofilm in mixed communities (figures S2, S4; table S4).

The second experiment aimed to mechanically disrupt the spatial structure of the community[20]. The original assay was performed with a third treatment that included swirling the colonies with a sterile metal sewing pin once a day. The results show only a slight, non-significant decrease in the fitness advantage of the biofilm strains (figure S5; table S5). We hypothesize that this result may be due to the frequency of mechanical disruption: 24 hours is enough time for the biofilm strain to segregate and grow before being disrupted again.

Taken together, these experiments suggest that the ability to form a biofilm provides a strong advantage in direct competition.

### Competitions between biofilm and non-biofilm strains without spatial structure

In order to verify that the results really were due to the spatial structure and not simply due to differing growth abilities in various conditions and community compositions, the same competitions were performed with no structure at all— in well-mixed liquid culture (figure S6). The results of the homogenous community treatment (strains grown alone and subsequently compared in randomly assigned pairs), recapitulated the results from the agar plates with the change in biofilm frequency around 1. In contrast, the mixed community treatment, in which two strains were grown together, showed an overall *disadvantage* to biofilm-forming strains in direct competition. We hypothesize that this is due to the cost of producing the components of a biofilm without the associated benefits of spatial structure. These results support published findings that showed biofilm-forming strains derived from a single genetic background grew more slowly than their smooth counterparts in liquid culture, while the diameter of the complex colonies grew more quickly than that of the smooth colonies on agar surfaces [54].

### Competitions between biofilm and toxin-producing strains

Given the potential for yeast biofilms to gain a competitive advantage through their spatial use, and the known ability of yeast killer toxins to kill nearby sensitive cells, we sought to determine whether biofilm production protected cooperative cells or if active toxin was effective against cells enmeshed in a biofilm. Pairs of biofilm and toxin strains (K2 and K28) were competed against one another in both homogenous and mixed communities, with and without inducing biofilm formation, and with and without active toxin. The toxin-encoding viruses can be lost when strains are cultured at high temperature; therefore it was not possible to transform and fluorescently mark the toxin strains.

Both biofilm-forming strains were sensitive to the toxins, as determined by halo assays (figure S7); however, the toxins had different effects when grown in a mixed biofilm community. In direct and indirect competition with the K2 strain, when the toxin was not active (blue and yellow in figures 2a, S8), the biofilm strain was more fit in all treatments (table S6). Similar to competitions with other smooth strains, the strongest fitness benefit to biofilm formation was in mixed communities. In contrast, when the toxin was active (orange and purple) in mixed communities, biofilm-forming cells were susceptible to the toxin, as indicated by the strong decrease in biofilm strain frequency. Inspection of the images shows the toxin strain mostly surrounding the biofilm and dominating the edge of the colony (figure 2c, videos S1-S3). However, in most cases, the increased spatial use by the biofilm allowed an escape at the edge of at least one section of the colony (arrows in figure 2c).

**Figure 2:**
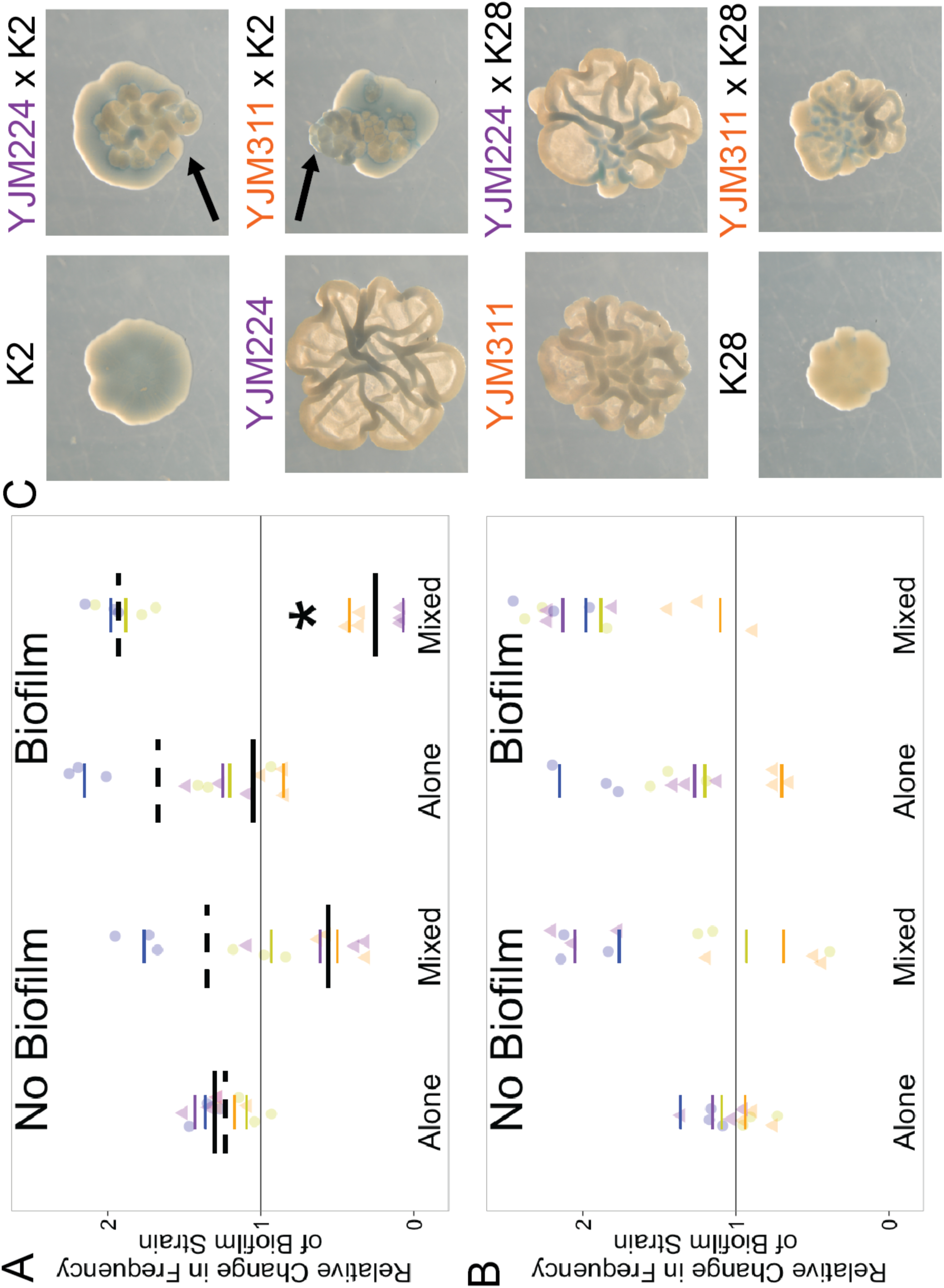
Fitness Effects of Biofilm Formation in the Presence of Killer Toxin. Biofilm-forming strains were competed against toxin-producing strains in pure and mixed colonies, with and without inducing biofilm formation, and with and without active toxin; a total of 144 colonies were assayed. Yellow and blue circles correspond to fitness assays of YJM311 and YJM224, respectively, against a toxin strain, but without active toxin (as in Figure 1; toxin strain is simply another environmental isolate); dashed line indicates overall mean.Orange and purple triangles refer to the same competitions, but with active toxin (low pH versions of the media); solid line indicates overall mean. **(A)** Assay with a K2 toxin-producing strain. * indicates significance at p< 0.0001. **(B)** Assay with a K28 toxin-producing strain. In mixed communities, YJM311 is significantly more fit without K28 toxin than with K28 toxin (*p*< 0.0001); however, YJM311 is more fit in the presence of K28 toxin when biofilms are induced compared to when they are not (*p*=0.03). **(C)** Representative images of single strain (left column) and mixed colonies (right column) grown on medium in which toxin is active and biofilm formation is induced (low pH, LD). Arrows indicate location of an escape of the biofilm strain at the edge of the colony. Blue dye indicates cell death.

Similarly, in direct and indirect competition with the K28 strain, when the toxin was not active (blue and yellow in figure 2b, S8), the biofilm strain was more fit in all treatments (table S7). However, in contrast to K2, active K28 was not effective in containing the biofilm strain. YJM224 was particularly resistant to the toxin, although there is cell death, as indicated by the presence of blue dye within the colonies. The growth of YJM311 was clearly altered, but the biofilm was able to escape in all cases.

Our results provide insight into a relationship between natural phenotypes that had not yet been explored: at least one killer toxin is effective against cells enmeshed in a biofilm, but biofilm formation may allow a sensitive strain a spatial escape.

## Discussion

This study investigated the fitness effects of biofilm formation in environmental isolates of the model organism, *Saccharomyces cerevisiae*; our results suggest a robust advantage in direct competition with non-biofilm-formers in spatially structured communities. In mixed colonies with biofilms induced (and without active toxin), biofilm strains consistently increased in frequency. Our results support the findings of bacterial and computational studies that show a competitive advantage associated with adhesion and spatial structure [10, 20-23], suggesting that eukaryotic microbial systems may function in a similar way.

Previous theoretical work has shown that in expanding smooth colonies containing two genotypes, founder effects lead to sectors (as seen in figure 1*b*); straight lines separating the boundaries of the sectors suggest a lack of competitive advantage, while curves suggest competition between the genotypes [56]. The mixed smooth colonies in Figures 1*b* and S2 suggest that the different environmental backgrounds compete with one another. This supports the idea that there is also competition in mixed colonies with biofilm formation, and that biofilms may serve a competitive function.

In contrast to the mixed strain colonies, our data did not show a fitness advantage to biofilm formation in indirect competition between single-strain colonies. These results are in agreement with Regenberg et al. (2016), who showed that the fitness benefit of yeast biofilm formation increased as the viscosity of the medium decreased; at 2% agar, the concentration used here, the dry biomass of biofilm and non-biofilm colonies was not different [55]. While *S. cerevisiae* has been isolated in numerous ecological niches, usually associated with fruits or man-made environments, it is unclear what habitat it evolved in and was historically adapted to [83]. Therefore, it is impossible to recapitulate natural conditions, making the choice of agar concentration somewhat arbitrary. Regardless, it is interesting that even in conditions that do not provide a fitness advantage to biofilm formation in indirect competition, biofilms still provide a competitive advantage in direct competition.

Next, this study investigated a less-understood interaction between two social phenotypes: biofilm formation and toxin production. Our results suggest that in this eukaryotic system, two toxins have different abilities in direct competition against biofilm-forming strains; K2 was effective at containing the growth of the biofilm strain, while K28 was not. The reason for this difference is unknown, but may be due to the different modes of action of the toxins. Even in the presence of the effective toxin, both biofilm strains were able to reach the leading edge of the colony and grow outward, generating a spatial escape. While the assay was performed in an artificial lab setting, the results suggest that increased use of space by yeast biofilms may not only provide an escape from competition for nutrients, but may also provide an escape from warfare phenotypes. It is notable that both the toxins and Flo11, the cellular adhesin responsible for cell-cell attachment, are most effective in acidic conditions[53, 78], thus suggesting that yeast biofilms and toxins likely interact in the environment.

## Conclusion

Biofilms represent the earliest form of multicellular structures in the evolution of life, and are currently found in numerous natural and man-made environments— from water filtration systems, to dental surfaces, to medical implants— and can pose a serious threat to human health. Thus, understanding biofilms not only leads to insights into the evolution of early microbial communities, but may have practical implications. Our study demonstrates that biofilms may provide similar benefits to eukaryotic lineages as they do to bacterial lineages, and suggests that eukaryotic microbes may meet the assumptions of much of microbial social evolution theory. Furthermore, we show that the premier biomedical model yeast, *S. cerevisiae*, may be a powerful system to investigate questions surrounding social evolution in eukaryotic biofilms, an area of research that has received little attention.

## Data

All data associated with the research are available in the electronic supplementary material.

## Competing Interests

We have no competing interests.

## Authors’ Contributions

ARH developed the biofilm assay and conducted experiments with environmental isolates; BMD conducted and designed experiments with environmental and toxin isolates, and participated in drafting the manuscript; BAL conducted liquid competitions; HAM designed experiments, analyzed data, and drafted the manuscript.

## Acknowledgements

We thank T. Meier and A. Reft for technical assistance; P. Magwene and D. Wloch-Salamon for strains; M. Pieczynska for advice on toxin assays.

## Funding

P41 RR11823 from the National Institutes of Health to T. N. Davis at UW for fluorescence cassettes, the WM Charles Center Dintersmith Fellowship to BMD, and the Jeffress Trust for Interdisciplinary Research award to HAM.

## References

1. West, S.A., et al., The Social Lives of Microbes. Annual Review of Ecology, Evolution and Systematics, 2007. 38: p. 53–77.

2. O'Toole, G., H.B. Kaplan, and R. Kolter, Biofilm formation as microbial development. Annual Review of Microbiology, 2000. 54(1): p. 49–79.

3. Lee, K.W.K., et al., Biofilm development and enhanced stress resistance of a model, mixed-species community biofilm. ISME J, 2014. 8(4): p. 894–907.

4. Nutman, A.P., et al., Rapid emergence of life shown by discovery of 3,700-million-year-old microbial structures. Nature, 2016. 537(7621): p. 535–538.

5. West, S.A., et al., Social evolution theory for microorganisms. Nat Rev Micro, 2006. 4(8): p. 597–607.

6. Crespi, B.J., The evolution of social behavior in microorganisms. Trends in Ecology and Evolution, 2001. 16(4): p. 178–183.

7. Rainey, P.B. and K. Rainey, Evolution of cooperation and conflict in experimental bacterial populations. Nature, 2003. 425(6953): p. 72–4.

8. Brockhurst, M.A., A. Buckling, and A. Gardner, Cooperation Peaks at Intermediate Disturbance. Current Biology, 2007. 17(9): p. 761–765.

9. Hallatschek, O., et al., Genetic drift at expanding frontiers promotes gene segregation. Proceedings of the National Academy of Sciences, 2007. 104(50): p. 19926–19930.

10. Xavier, J.B. and K.R. Foster, Cooperation and conflict in microbial biofilms. Proceedings of the National Academy of Sciences, 2007. 104(3): p. 876–881.

11. Nadell, C.D., K.R. Foster, and J.B. Xavier, Emergence of Spatial Structure in Cell Groups and the Evolution of Cooperation. PLOS Computational Biology, 2010. 6(3): p. e1000716.

12. Nadell, C.D. and B.L. Bassler, A fitness trade-off between local competition and dispersal in Vibrio cholerae biofilms. Proceedings of the National Academy of Sciences, 2011. 108(34): p. 14181–14185.

13. Momeni, B., et al., Strong inter-population cooperation leads to partner intermixing in microbial communities. eLife, 2013. 2: p. e00230.

14. Anderson, M.S., E.C. Garcia, and P.A. Cotter, Kind Discrimination and Competitive Exclusion Mediated by Contact-Dependent Growth Inhibition Systems Shape Biofilm Community Structure. PLOS Pathogens, 2014. 10(4): p. e1004076.

15. Millet, Y.A., et al., Insights into Vibrio cholerae Intestinal Colonization from Monitoring Fluorescently Labeled Bacteria. PLOS Pathogens, 2014. 10(10): p. e1004405.

16. Müller, M.J.I., et al., Genetic drift opposes mutualism during spatial population expansion. Proceedings of the National Academy of Sciences, 2014. 111(3): p. 1037–1042.

17. van Gestel, J., et al., Density of founder cells affects spatial pattern formation and cooperation in Bacillus subtilis biofilms. Isme j, 2014. 8(10): p. 2069–79.

18. Van Dyken, J.D., et al., Spatial Population Expansion Promotes the Evolution of Cooperation in an Experimental Prisoner's Dilemma. Current Biology. 23(10): p. 919–923.

19. Nadell, C.D., K. Drescher, and K.R. Foster, Spatial structure, cooperation and competition in biofilms. Nat Rev Micro, 2016. 14(9): p. 589–600.

20. Kim, W., et al., Importance of positioning for microbial evolution. Proc Natl Acad Sci U S A, 2014. 111(16): p. E1639–47.

21. Irie, Y., et al., The Pseudomonas aeruginosa PSL Polysaccharide Is a Social but Noncheatable Trait in Biofilms. MBio, 2017. 8(3).

22. Garcia, T., G. Doulcier, and S. De Monte, The evolution of adhesiveness as a social adaptation. Elife, 2015. 4.

23. Schluter, J., et al., Adhesion as a weapon in microbial competition. ISME J, 2015. 9(1): p. 139–149.

24. Oliveira, N.M., et al., Biofilm Formation As a Response to Ecological Competition. PLOS Biology, 2015. 13(7): p. e1002191.

25. Riley, M.A. and J.E. Wertz, Bacteriocins: evolution, ecology, and application. Annu Rev Microbiol, 2002. 56: p. 117–37.

26. Schmitt, M.J. and F. Breinig, Yeast viral killer toxins: lethality and self-protection. Nat Rev Microbiol, 2006. 4(3): p. 212–21.

27. Tait, K. and I.W. Sutherland, Antagonistic interactions amongst bacteriocin-producing enteric bacteria in dual species biofilms. J Appl Microbiol, 2002. 93(2): p. 345–52.

28. Gardner, A. and S.A. West, Spite and the scale of competition. Journal of Evolutionary Biology, 2004. 17(6): p. 1195–1203.

29. Bucci, V., C.D. Nadell, and J.B. Xavier, The evolution of bacteriocin production in bacterial biofilms. Am Nat, 2011. 178(6): p. E162–73.

30. Weber, M.F., et al., Chemical warfare and survival strategies in bacterial range expansions. Journal of The Royal Society Interface, 2014. 11(96).

31. Abrudan, M.I., et al., Socially mediated induction and suppression of antibiosis during bacterial coexistence. Proceedings of the National Academy of Sciences, 2015. 112(35): p. 11054–11059.

32. McNally, L., et al., Killing by Type VI secretion drives genetic phase separation and correlates with increased cooperation. Nat Commun, 2017. 8: p. 14371.

33. West, S.A. and G.A. Cooper, Division of labour in microorganisms: an evolutionary perspective. Nat Rev Microbiol, 2016. 14(11): p. 716–723.

34. Blankenship, J.R. and A.P. Mitchell, How to build a biofilm: a fungal perspective. Current Opinion in Microbiology, 2006. 9(6): p. 588–594.

35. Nobile, C.J. and A.D. Johnson, Candida albicans Biofilms and Human Disease, in Annual Review of Microbiology, Vol 69, S. Gottesman, Editor. 2015. p. 71–92.

36. Kuthan, M., et al., Domestication of wild Saccharomyces cerevisiae is accomplanied by changes in gene expression and colony morphology. Molecular Microbiology, 2003. 47(3): p. 745–754.

37. Reynolds, T.B. and G.R. Fink, Bakers' Yeast, a Model for Fungal Biofilm Formation. Science, 2001. 291(5505): p. 878–881.

38. Verstrepen, K.J. and F.M. Klis, Flocculation, adhesion and biofilm formation in yeasts. Molecular Microbiology, 2006. 60(1): p. 5–15.

39. Honigberg, S.M., Cell signals, cell contacts, and the organization of yeast communities. Eukaryotic Cell, 2011. 10(4): p. 466–473.

40. Vallejo, J., et al., Cell aggregations in yeasts and their applications. Applied Microbiology and Biotechnology, 2013. 97(6): p. 2305–2318.

41. Liti, G., et al., Population genomics of domestic and wild yeasts. Nature, 2009. 458: p. 337–341.

42. Strope, P.K., et al., The 100-genomes strains, an S. cerevisiae resource that illuminates its natural phenotypic and genotypic variation and emergence as an opportunistic pathogen. Genome Research, 2015. 25(5): p. 762–774.

43. Schacherer, J., et al., Comprehensive polymorphism survey elucidates population sturcture of Saccharomyces cerevisiae. Nature, 2009. 458: p. 342–346.

44. Cromie, G.A., et al., Genomic Sequence Diversity and Population Structure of Saccharomyces cerevisiae Assessed by RAD-seq. G3: Genes Genomes Genetics, 2013. 3(12): p. 2163–2171.

45. Ludlow, Catherine L., et al., Independent Origins of Yeast Associated with Coffee and Cacao Fermentation. Current Biology, 2016. 26(7): p. 965–971.

46. Hope, E.A. and M.J. Dunham, Ploidy-Regulated Variation in Biofilm-Related Phenotypes in Natural Isolates of Saccharomyces cerevisiae. G3: Genes Genomes Genetics, 2014. 4(9): p. 1773–1786.

47. Pieczynska, M.D., J.A.G.M. de Visser, and R. Korona, Incidence of symbiotic dsRNA 'killer' viruses in wild and domesticated yeast. FEMS Yeast Research, 2013. 13(8): p. 856–859.

48. Granek, J.A. and P.M. Magwene, Environmental and Genetic Determinants of Colony Morphology in Yeast. PLoS Genet, 2010. 6(1): p. e1000823.

49. Brückner, S. and H.-U. Mösch, Choosing the right lifestyle: adhesion and development in Saccharomyces cerevisiae. FEMS Microbiological Review, 2011. 36: p. 25–58.

50. Bojsen, R.K., K.S. Andersen, and B. Regenberg, *Saccharomyces cerevisiae* – a model to uncover molecular mechanisms for yeast biofilm biology. FEMS Immunology & Medical Microbiology, 2012. 65(2): p. 169–182.

51. Štovíček, V., et al., General factors important for the formation of structured biofilm-like yeast colonies. Fungal Genetics and Biology, 2010. 47: p. 1012–1022.

52. Váchová, L., et al., Flo11p, drug efflux pumps, and the extracellular matrix cooperate to form biofilm yeast colonies. The Journal of Cell Biology, 2011. 194(5): p. 679–687.

53. Kraushaar, T., et al., Interactions by the Fungal Flo11 Adhesin Depend on a Fibronectin Type III-like Adhesin Domain Girdled by Aromatic Bands. Structure. 23(6): p. 1005–1017.

54. Tan, Z., et al., Aneuploidy underlies a multicellular phenotypic switch. Proceedings of the National Academy of Sciences, 2013. 110(30): p. 12367–12372.

55. Regenberg, B., et al., Clonal yeast biofilms can reap competitive advantages thorough cell differentiation without being obligatorily multicellular. Proceedings of the Royal Society B-Biological Sciences, 2016. 283.

56. Korolev, K.S., et al., Selective sweeps in growing microbial colonies. Phys Biol, 2012. 9(2): p. 026008.

57. Chen, L., et al., Two-Dimensionality of Yeast Colony Expansion Accompanied by Pattern Formation. Plos Computational Biology, 2014. 10(12).

58. Orentaite, I., et al., K2 killer toxin-induced physiological changes in the yeast Saccharomyces cerevisiae. FEMS Yeast Res, 2016. 16(2): p. fow003.

59. Serviene, E., et al., Screening the Budding Yeast Genome Reveals Unique Factors Affecting K2 Toxin Susceptibility. PLoS ONE, 2012. 7(12): p. e50779.

60. Novotna, D., H. Flegelova, and B. Janderova, Different action of killer toxins K1 and K2 on the plasma membrane and the cell wall of Saccharomyces cerevisiae. FEMS Yeast Res, 2004. 4(8): p. 803–13.

61. Wingfield, B.D., et al., K3 killer yeast is a mutant K2 killer yeast. Mycological Research, 1990. 94(7): p. 901–6.

62. Pfeiffer, P. and F. Radler, Purification and Characterization of Extracellular and Intracellular Killer Toxin of Saccharomyces cerevisiae Strain 28. Microbiology, 1982. 128(11): p. 2699–2706.

63. Schmitt, M.J., et al., Cell cycle studies on the mode of action of yeast K28 killer toxin. Microbiology, 1996. 142 (Pt 9): p. 2655-62.

64. Kreft, J.U., Biofilms promote altruism. Microbiology, 2004. 150(Pt 8): p. 2751-60.

65. Reuter, M., G. Bell, and D. Greig, Increased outbreeding in yeast in response to dispersal by an insect vector. Current Biology, 2007. 17(3): p. R81–R83.

66. Stefanini, I., et al., Role of social wasps in Saccharomyces cerevisiae ecology and evolution. Proceedings of the National Academy of Sciences, 2012. 109(33): p. 13398–13403.

67. Stefanini, I., et al., Social wasps are a Saccharomyces mating nest. Proceedings of the National Academy of Sciences, 2016. 113(8): p. 2247–2251.

68. Goddard, M.R., et al., A distinct population of *Saccharomyces cerevisiae* in New Zealand: evidence for local dispersal by insects and human-aided global dispersal in oak barrels. Environmental Microbiology, 2009. 12(1): p. 63–73.

69. McCusker, J.H., et al., Genetic characterization of pathogenic Saccharomyces isolates. Genetics, 1994. 136: p. 1261–1269.

70. McCullough, M.J., et al., Epidemiological investigation of vaginal Saccharomyces cerevisiae isolates by a genotypic method. J Clin Microbiol, 1998. 36(2): p. 557–62.

71. Sniegowski, P.D., P.G. Dombrowski, and E. Fingerman, *Saccharomyces cerevisiae* and Saccharomyces paradoxus coexist in a natural woodland site in North America and display diffferent levels of reproductive isolation from European conspecifics. FEMS Yeast Research, 2002. 1: p. 299–306.

72. Gietz, R. and R. Woods, Transformation of yeast by the Liac/Ss Carrier Dna/Peg method. Methods in Enzymology, 2002. 350: p. 87–96.

73. Wach, A., et al., New heterologous modules for classical or PCR-based gene disruptions in *Saccharomyces cerevisiae*. Yeast, 1994. 10: p. 1793–1808.

74. Goldstein, A. and J. McCusker, Three new dominant drug resistance cassettes for gene disruption in Saccharomyces cerevisiae. Yeast, 1999. 15: p. 1541–1553.

75. Longtine, M.S., et al., Additional modules for versatile and economical PCR-based gene deletion and modification in Saccharomyces cerevisiae. Yeast, 1998. 14(10): p. 953–61.

76. Hailey, D.W., T.N. Davis, and E.G. Muller, Fluorescence resonance energy transfer using color variants of green fluorescent protein. Methods Enzymol, 2002. 351: p. 34–49.

77. Schmitt, M.J. and D.J. Tipper, K28, a unique double-stranded RNA killer virus of Saccharomyces cerevisiae. Mol Cell Biol, 1990. 10(9): p. 4807–15.

78. Lukša, J., S. Serva, and E. Servienė, *Saccharomyces cerevisiae* K2 toxin requires acidic environment for unidirectional folding into active state. Mycoscience, 2016. 57: p. 51–57.

79. Suzuki, Y., et al., Cysteine residues in a yeast viral A/B toxin crucially control host cell killing via pH-triggered disulfide rearrangements. Mol Biol Cell, 2017. 28(8): p. 1123–1131.

80. Woods, D.R. and E.A. Bevan, Studies on the nature of the killer factor produced by Saccharomyces cerevisiae. Journal of General Microbiology, 1968. 51: p. 115–126.

81. Schindelin, J., et al., Fiji: an open-source platform for biological-image analysis. Nat Methods, 2012. 9(7): p. 676–82.

82. Chevin, L.-M., On measuring selection in experimental evolution. Biology Letters, 2010.

83. Goddard, M.R. and D. Greig, Saccharomyces cerevisiae: a nomadic yeast with no niche? FEMS Yeast Research, 2015. 15(3): p. fov009–fov009.

